# Widespread intronic polyadenylation diversifies immune cell transcriptomes

**DOI:** 10.1101/158998

**Authors:** Irtisha Singh, Shih-Han Lee, Mehmet K Samur, Yu-Tzu Tai, Mehmet K Samur, Yu-Tzu Tai, Nikhil C Munshi, Christine Mayr, Christina S. Leslie

## Abstract

Alternative cleavage and polyadenylation (ApA) can generate mRNA isoforms with differences in 3’UTR length without changing the coding region (CDR). However, ApA can also recognize intronic polyadenylation (IpA) signals to generate transcripts that lose part or all of the CDR. We analyzed 46 3’-seq and RNA-seq profiles from normal human tissues, primary immune cells, and multiple myeloma (MM) samples and created an atlas of 4,927 high confidence IpA events. Up to 16% of expressed genes in immune cells generate IpA isoforms, a majority of which are differentially used during B cell development or in different cellular environments, while MM cells display a striking loss of IpA isoforms expressed in plasma cells, their cell type of origin. IpA events can lead to truncated proteins lacking C-terminal functional domains. This can mimic ectodomain shedding through loss of transmembrane domains or alter the binding specificity of proteins with DNA-binding or protein-protein interaction domains, thus contributing to diversification of the transcriptome.

## Introduction

ApA is generally viewed as the selection of ApA signals in the 3’UTR, leading to the expression of different 3’UTR isoforms that code for the same protein. Recent computational analyses of 3’-end sequencing data have characterized the nature and extent of ApA in mammalian 3’UTRs^1,2,3,4,5,6,7^. For example, analysis of the first human ApA tissue atlas established that half of human genes express multiple 3’UTRs, enabling tissue-specific post-transcriptional regulation of ubiquitously expressed genes^1^. However, ApA events can also occur in introns rather than 3’UTRs, generating either non-coding transcripts or transcripts with truncated coding regions that lead to loss of C-terminal domains in the protein product.

The most famous example of cell-type specific usage of an IpA signal occurs in the immunoglobulin M heavy chain *(IGHM)* locus^8,9^. In mature B cells, recognition of the polyA signal in the 3’UTR produces the full-length message, including two terminal exons that encode the transmembrane domain of the plasma membrane-bound form of IgM (Fig. 1a). In plasma cells, usage of an IpA signal instead results in expression of an IpA isoform lacking these two terminal exons, leading to loss of the transmembrane domain and secretion of IgM antibody. Many additional IpA-generated truncated proteins have been described,^10,11^ including the soluble forms of EGF and FGF receptors and a truncated version of the transcription factor NFI-B^12^. One of the first examples of IpA was the generation of two isoforms of the interferon-induced anti-viral enzyme OAS1^13^. Whereas the IpA and the full-length mRNA transcripts encode enzymes with comparable enzymatic activity, the shorter transcript generates a hydrophobic C-terminus and the longer transcript an acidic C-terminus. This suggests that the two isoforms may interact with different co-factors or cellular structures^13^. Other examples include the transcription factor SREPF, whose IpA isoform can act as a developmental switch during spermatogenesis^14^.

**Figure 1:**
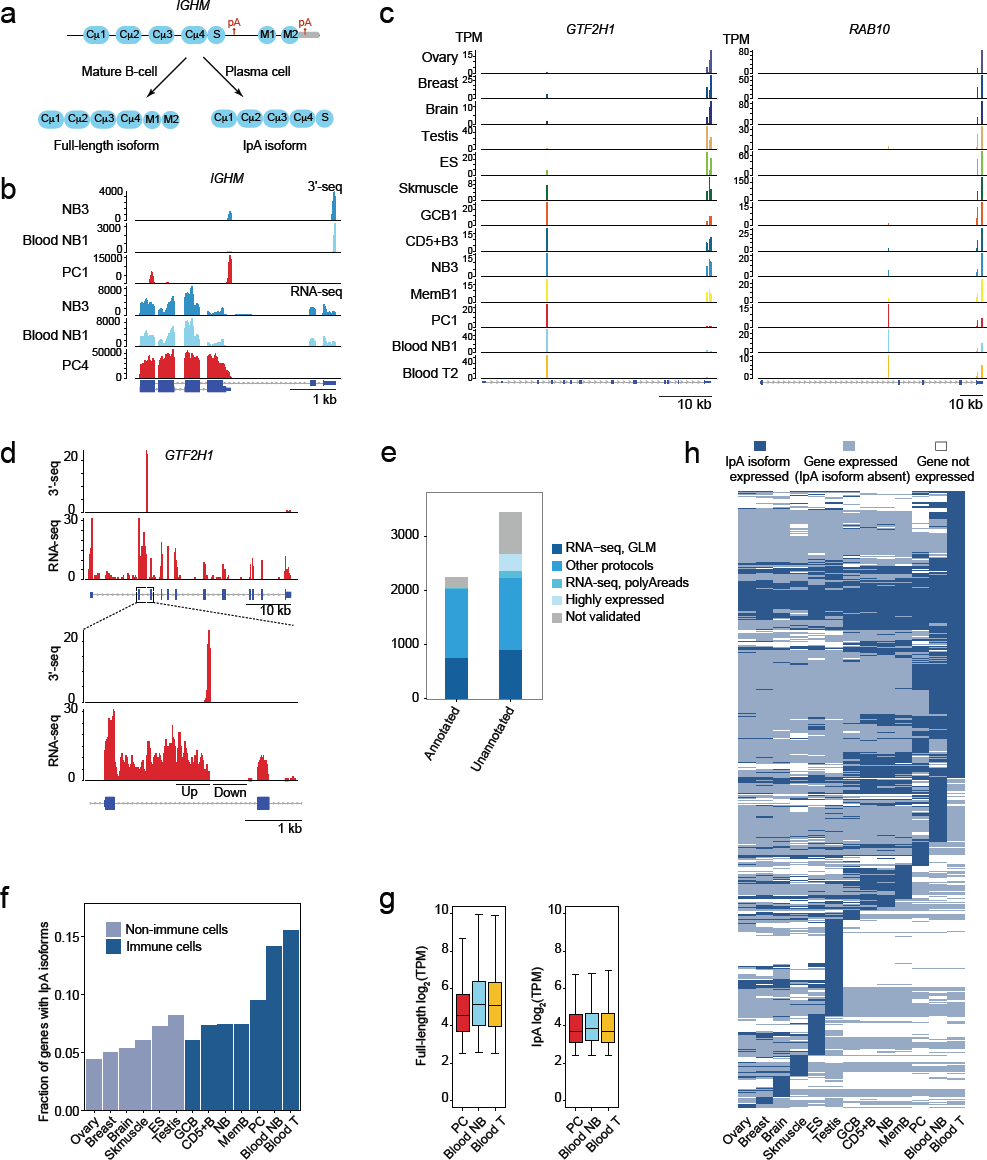
Widespread intronic polyadenylation with robust expression in circulating immune cells. a)Schematic representation of full-length and IpA isoform of *IGHM* expressed in mature B cells and plasma cells (PC).b)3’-seq (tags per million, TPM) and RNA-seq (read coverage) tracks showing expression of the IpA and full-length mRNA isoforms of *IGHM* (ENSG00000211899), encoding the immunoglobulin mu heavy chain, IgM. The full-length isoform is expressed in NB from blood and lymphoid tissue and includes two exons encoding the C-terminal transmembrane domain of membrane-bound IgM. The IpA isoform is expressed in PCs obtained from bone marrow. It lacks the transmembrane domain which leads to expression of soluble IgM.c)3’-seq tracks showing IpA isoform expression for two genes across human tissues and immune cell types.d)RNA-seq coverage of intronic regions flanking IpA sites. A GLM-based test is used to validate the IpA isoforms. An isoform is considered validated if there is a significant difference (FDR-adjusted *p* < 0.1) in read counts in windows located up-and downstream of the putative IpA site.e)The fraction of IpA isoforms validated by read evidence from independent data sets is shown for annotated and unannotated IpA isoforms. IpA isoforms present in RefSeq, UCSC genes and Ensembl databases are considered to be annotated.f)The fraction of expressed genes that generate IpA isoforms is shown for each cell type.g)Expression levels (log_2_ TPM) for full-length mRNAs and IpA isoforms are shown as boxplots for in PCs, blood NB and T cells. IpA isoforms are robustly expressed as full-length mRNA expression is 4.53, 5.15 and 5.11, respectively, compared to 3.71, 3.83 and 3.71 for IpA isoforms.h)Tissue-specific expression of IpA isoforms. Each row represents a gene. Dark blue, IpA isoform is expressed (> 5 TPM); light blue, IpA isoform is not expressed, but full-length mRNA is expressed (> 5.5 TPM); and white, gene is not expressed.

In the splicing literature, isoforms generated through recognition of an IpA signal are often described as ‘alternative last exon’ events^15^. It is thought that genes that generate IpA isoforms harbor competing splicing and polyA signals. When splicing outcompetes polyadenylation, a full-length mRNA is generated, and otherwise a truncated mRNA is made^16^. As the defining event is the recognition of an IpA signal, we call these transcripts IpA isoforms. Only through the analysis of 3’ end sequencing data has it been possible to recognize the widespread expression of IpA isoforms, and here we present a systematic analysis of IpA isoforms across diverse cell types.

We adapted our previous computational analysis pipeline for 3’-seq to identify robust ApA events that occur in introns and to quantify expression of IpA isoforms across human tissues, immune cells, and in MM patient samples. We focused on immune cells because it is feasible to obtain pure cell populations of primary cells and because in our previous tissue atlas we found B cells to express the largest number of differential 3’UTR isoforms^1^. Through integration with RNA-seq profiles in B lineage and MM cells as well as external data sets and annotation databases, we assembled an atlas of confident IpA isoforms that are either supported by independent data sources or are very highly expressed in at least one cell type in our data set. We found that IpA isoforms are widely expressed, most prevalently in blood-derived immune cells, and that generation of IpA isoforms is regulated during B cell development, between cellular environments and in cancer. IpA events in immune cells are enriched at the start of the transcription unit, leading to IpA isoforms that retain none or little of the CDR and hence represent a novel class of robustly expressed non-coding transcripts. The majority of IpA events that occur later in transcription units can lead to truncated proteins often lacking repeated C-terminal functional domain, and thus contribute to the diversification of the transcriptome.

## Results

### Computational analysis of 3’-seq reveals widespread intronic polyadenylation

To assemble an atlas of IpA isoforms in human tissues and immune cells, we used our previously published 3’-seq data set from normal human tissues (ovary, brain, breast, skeletal muscle, testis), cell types (embryonic stem (ES) cells, naïve B cells from peripheral blood (blood NB)), and cell lines^1^ and combined it with a newly generated data set from normal and malignant primary immune cells. The new immune cell profiles (n = 29) were all performed with biological replicates and included lymphoid tissue-derived naïve B cells (NB), memory B cells (MemB), germinal center B cells (GCB) and CD5+ B cells (CD5+B), blood T cells and plasma cells (PC) and MM derived from bone marrow aspirates (**Supplementary Tables 1, 2**). We adapted our previously described computational pipeline to process 3’-seq libraries and detect and quantify ApA events, including intronic as well as 3’UTR events, while removing technical artifacts (see **Methods**)^1^. All subsequent analyses were restricted to protein coding genes. For additional evidence in support of IpA isoforms, we performed RNA-seq profiling in the same normal and malignant B cell types, where possible for the same samples (**Supplementary Table 3**).

We confirmed from both 3’-seq and RNA-seq data that the IpA isoform of *IGHM* is highly expressed in PC while the full-length transcript, encoding membrane-bound IgM, is the dominant isoform in NB cells (Fig. 1b). Analysis of 3’-seq also revealed novel putative IpA isoforms, including in the locus of *GTF2H1,* encoding a subunit of general transcription factor II H, and *RAB10,* encoding a member of the Ras oncogene family of small GTPases (Fig. 1c). Like 3’UTR isoforms, IpA isoforms display differential expression across tissues and cell types. For example, the IpA isoform of *GTF2H1* is well expressed in skeletal muscle and all immune cell types assayed, and indeed the IpA isoform is the only isoform expressed from this gene in PC, blood NB and T cells; these three cell types are also the only ones to express the IpA isoform of *RAB10.* To validate transcriptome-wide the IpA events identified by 3’-seq, we used RNA-seq data from the same cell types. We expect intronic read coverage upstream but not downstream of the IpA event, as is visible in RNA-seq coverage in PCs flanking the intronic 3’-seq peak in *GTF2H1* (Fig. 1d). Formally, we can test if RNA-seq read counts are significantly higher in intronic windows chosen upstream compared to downstream of IpA events (see **Methods**)^17^. Significantly differential coverage could be confirmed for 29% (n = 1,670) of IpA events from our 3’-seq peak calls (FDR-adjusted *p* < 0.1), whereas almost no significant read count differences were found relative to randomly chosen positions in introns (**Supplementary Fig. 1a**). To assemble an atlas of highly confident IpA events for further analysis, we compared each intronic peak detected in the 3’-seq data against external annotation and data sources to find additional evidence in support of the IpA isoform (see **Methods, Supplementary Fig. 1b, c**). Briefly, IpA events that overlapped with the last exon of annotated isoforms in RefSeq, UCSC and Ensembl were first added to the atlas (2,241 events); unannotated IpA events that satisfied the test for differential upstream vs. downstream RNA-seq coverage were added next (907 events); unannotated IpA events without differential RNA-seq coverage but supported in data sets from other 3’ end sequencing protocols were then added to the atlas (1,332 events)^18^. We then added IpA events that lacked the previous sources of evidence but had RNA-seq support of the cleavage event – i.e. reads overlapping untemplated adenosines in the polyA tail (124 events). Finally, events with high expression in at least one cell type were also included in the atlas (323 events). 13% (n = 743) of IpA events could not be validated by any of these methods and thus were not included in the atlas for further analysis (**Supplementary Fig. 1c**). Overall, the atlas contains 4,927 confident IpA events in 3,431 protein coding genes, and 55% of atlas events are unannotated in RefSeq, UCSC and Ensembl (**Supplementary Fig. 1c**). We also found that similar proportions of annotated and unannotated IpA isoforms were validated by various kinds of supporting evidence (Fig. 1e); for example, only a slightly larger fraction of annotated vs. unannotated IpA events are supported by RNA-seq coverage (34% vs. 26%).

### IpA isoforms are robustly expressed in circulating immune cells

Using our atlas of IpA isoforms, we determined the prevalence of IpA across normal tissues and cell types by computing the fraction of genes expressing at least one IpA isoform out of all expressed genes in each cell type (Fig. 1f). Blood T cells had the highest fraction of genes with IpA isoforms (0.16) while ovary had the lowest fraction (0.04). Whereas 6-16% of genes expressed in immune cells generate IpA isoforms, complex tissues produce IpA isoforms in only 4-8% genes. Notably, blood NB cells expressed 1,114 IpA isoforms compared to only 721 IpA isoforms for tissue-derived NB cells, suggesting that the cellular environment has a strong effect on IpA isoform expression.

IpA isoforms are robustly expressed, with median expression of the same order of magnitude as the median expression of full-length isoforms (log_2_TPM of 3.71-3.83 for IpA isoforms in PC, blood NB, and blood T as compared to log_2_ TPM of 4.53-5.15 for full-length transcripts; Fig. 1g). Therefore, IpA isoforms are not ‘transcriptional noise’ produced from recognition of ‘cryptic’ sites, but rather represent major mRNA isoforms generated from alternative mRNA processing.

Figure 1h shows the tissue-specific expression of IpA isoforms. The mean expression level of the IpA isoform across the replicates had to be more than 5 TPM to be considered as an expressed IpA isoform. The heatmap shows that a majority of the IpA isoforms with reproducible expression patterns are expressed in immune cell types (n = 3,365), and almost all of these are expressed in at least two immune cell types. Non-immune tissues like testis and ES cells express tissue-specific IpA isoforms, but the majority of these isoforms are produced from tissue-specific genes.

### Cell types with frequent IpA retain less coding region in IpA isoforms and express shorter 3’UTRs

To begin to assess the impact of IpA isoform expression, we computed the fraction of retained CDR for each IpA isoform based on the position of the IpA event relative to the full-length annotated CDR. The histogram of retained CDR fraction across the IpA atlas showed a uniform distribution across the CDR except for a substantial overrepresentation of IpA isoforms that lose all or almost all of the CDR (Fig. 2a). However, an examination of histograms of retained CDR fraction across individual tissues and cell types revealed a more nuanced picture (Fig. 2b), where IpA events near the start of the transcription unit dominate in blood and bone marrow-derived immune cells, while brain and ES cells preferentially generate IpA events close to the end of transcription units. In testis and tissue-derived B cells, we found an intermediate pattern.

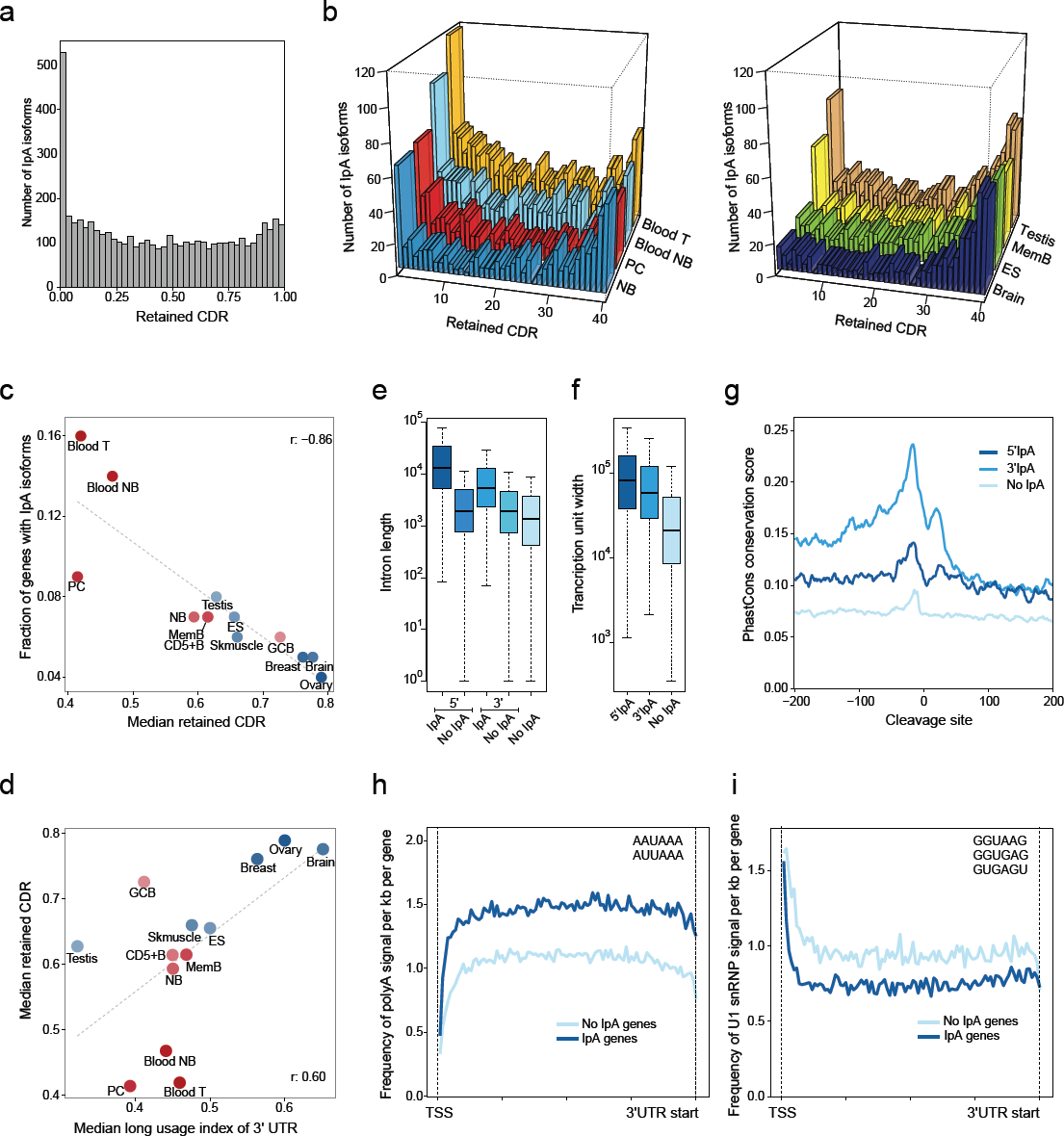
Enrichment of IpA sites at the start of transcription units. a)The fraction of retained coding region (CDR) was calculated as the nucleotides from the start codon to the end of the exon located upstream of the IpA peak, divided by all coding nucleotides of the longest annotated open reading frame and is shown for all IpA isoforms in the atlas.b)As in (a), but shown for individual cell types.c)Correlation between the median retained CDR with the fraction of genes that generate IpA isoforms in each sample (Pearson correlation coefficient, r = −0.86). Tissues with a higher proportion of IpA isoforms generate IpA isoforms with shorter CDRs.d)Correlation between the median retained CDR and the median usage of the distal ApA site in the 3’UTR (Pearson correlation coefficient, r = 0.60). Tissues with shorter 3’UTRs have IpA isoforms with shorter CDRs.e)IpA isoforms occur in long introns. The introns in which 5’IpA events occur are longer than the other introns of the same genes (one-sided Wilcoxon rank-sum test, *p* < 10^−20^). Similarly, the introns in which 3’IpA events occur are longer than the remaining introns of those genes (onesided Wilcoxon rank-sum test, *p* < 10^−20^). If taken together, then the introns in which IpA events occur are longer than the introns of the genes that only express full-length isoform (one-sided Wilcoxon rank-sum test, *p* < 10^−20^).f)IpA isoforms occur in genes with long transcription units. Genes that express IpA isoforms have longer transcription units compared to genes that only express full-length isoforms (onesided Wilcoxon rank-sum test, *p* < 10^−20^).g)Higher conservation around the cleavage sites of IpA isoforms. The plot shows PhastCons scores of 200 nt upstream and downstream of IpA cleavage sites (x = 0). 5’IpA and 3’IpA events both have significantly higher conservation flanking the cleavage site compared to corresponding regions of randomly selected polyA signals (AAUAAA) in introns lacking IpA events (one-sided Wilcoxon signed-rank test, *p* < 10^−68^ for both comparisons).h)Genes with IpA isoforms (n = 3,481) are enriched for polyA sites compared with genes that do no generate IpA isoforms (n = 12,092) (one-sided Wilcoxon signed-rank test, *p* < 10^−18^). The frequency of polyA sites was counted from the TSS to the beginning of the 3’UTR and is shown as the average number of signals per kb per gene.i)As in (h), but U1 binding sites are shown. IpA genes are depleted for U1 snRNP signals (onesided Wilcoxon signed-rank test, *p* < 10^−18^).

We observed a significant negative correlation across tissues between the frequency of IpA isoform expression and length of retained CDR (r = −0.86, Fig. 2c). We further observed that cell types with a tendency to produce longer 3’UTRs also prefer to produce IpA isoforms that are located at the 3’ ends of transcription units (r = 0.60, Fig. 2d).

We use the term 5’IpA for IpA isoforms that retain less than 25% of the CDR and 3’IpA for the remainder. Both 5’IpA and 3’IpA events occur in introns that are significantly longer than the introns from the same genes that contain no IpA events or from genes that only express full-length transcripts (one-sided Wilcoxon rank-sum test, *p* < 10^−20^ for all three comparisons, Fig. 2e)^19^. Similarly, 5’IpA and 3’IpA isoforms are expressed from significantly longer transcription units than genes that only express full-length transcripts (one-sided Wilcoxon rank-sum test, *p* < 10^−20^ for both comparisons, Fig. 2f). 3’IpA atlas events have higher conservation by PhastCons in the sequence surrounding the polyA signal compared to 5’IpA atlas events (one-sided Wilcoxon signed-rank test, *p* < 10^−66^; Fig. 2g); however, 5’IpA events still show higher conservation than randomly chosen intronic polyA signals with no 3’-seq coverage (one-sided Wilcoxon signed-rank test, *p* < 10^−68^; Fig. 2g)^20^.

Previously, U1 snRNP expression and the presence of U1 snRNP motifs early in the transcription unit were found to play a crucial role in preventing premature cleavage and polyadenylation^21,22^. Consistent with these observations, we found that genes that express IpA isoforms contain a higher frequency of polyA signals within their transcription unit and are depleted for U1 snRNP signals, as compared to genes that only express 3’UTR isoforms (Fig. 2h, i). Therefore, genomic architecture and sequence composition may facilitate IpA isoform expression.

### IpA isoform expression is associated with moderate downregulation of full-length mRNAs

Next we used a generalized linear model (GLM) approach to determine significant changes in the relative of expression of IpA isoforms compared to full-length transcripts (usage of IpA) across normal immune cells (see **Methods**)^1,17,23^. The usage of the majority of expressed IpA isoforms differed significantly when we compared NB cells from lymphoid tissue and blood T cells (950/1308, Fig. 3a, FDR-adjusted *p* < 0.05). Within the B cell lineage, we also found differential usage of IpA between cell types, with PC in particular showing strikingly increased usage of IpA sites compared to tissue-derived NB cells (Fig. 3a). However, surprisingly, when we compared NB cells from lymphoid tissue and blood, we found even more significant changes in IpA usage (720/1113) than we observed between different B cell types (Fig. 3a, b). This indicates that IpA isoform expression is not only cell type-differential but also highly dynamic, changing between different cellular environments. Genes with differential usage of IpA isoforms between immune cell types were most strongly enriched for annotations including zinc finger domains, bromodomains, and the ubiquitin-like conjugation pathway (Fig. 3c)^24^.

**Figure 3:**
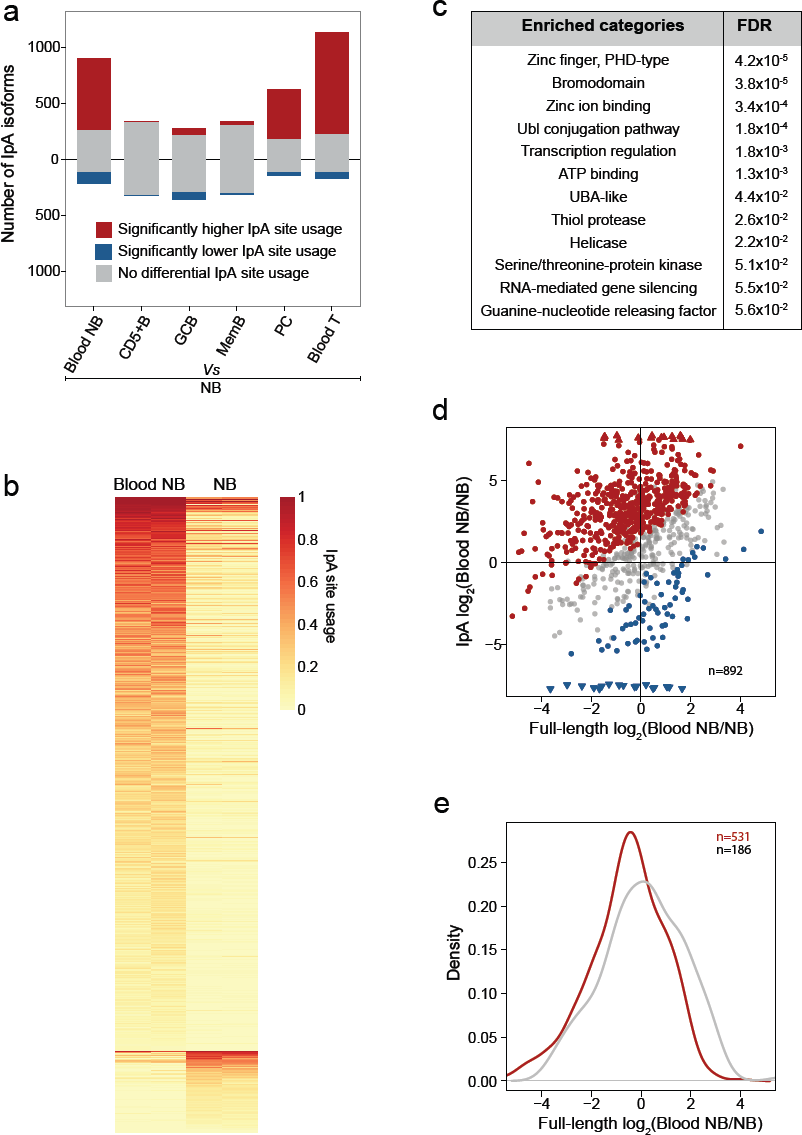
Dynamic expression of IpA isoforms in immune cells. a)Number of IpA isoforms with differential usage of IpA sites between NB from lymphoid tissue versus other immune cells (FDR-adjusted *p* < 0.05).b)Heatmap showing IpA site usage of IpA isoforms with significantly different usage (FDR-adjusted *p* < 0.05) between NB derived from blood or lymphoid tissue (n = 720). Each row indicates an IpA isoform.c)Enrichment of gene ontology terms for the genes shown in (b).d)Fold change of IpA isoform and full-length mRNA expression in blood versus lymphoid tissue-derived NB by TPM. All the genes that were tested for differential usage are shown (n = 892). If a gene had multiple IpA isoforms, then the one with the most significant differential IpA usage is shown. IpA isoforms with significantly different usage (FDR-adjusted *p* < 0.05) are highlighted in red (higher usage) or blue (lower usage).e)Significant downregulation of full-length mRNAs in genes with significant IpA isoform expression (one sided KS test, *p* < 10^−5^). Shown are genes highlighted in red from (d).

We then asked if cells might use IpA signals in a switch-like fashion to ‘turn off’ expression of the full-length transcript by ‘turning on’ the expression of IpA isoform. In Fig. 3d, we plot the expression change of the IpA isoform against that of the full-length transcript and show IpA genes that differentially increase (red points) or decrease (blue points) usage of their IpA isoforms in blood-derived compared with tissue-derived NB cells. If a gene had multiple IpA isoforms, then the one with the most significant differential IpA usage is shown. Genes that increase the usage of the IpA isoform in blood-versus tissue-derived NB cells significantly reduced expression of their full-length transcripts compared to genes without significant change in IpA usage (Fig. 3e, one sided KS test, *p* < 10^−5^). The decrease in full-length isoform expression was modest, but significant, indicating that IpA usage does not predominantly result in a ‘switch-like’ change between full-length and IpA isoform expression.

### IpA diversifies the transcriptome through loss of C-terminal functional domains

Next, we investigated the potential functional consequences of IpA. We observed that IpA genes encode full-length proteins that are significantly larger and contain more domains than genes that do not produce IpA isoforms (Fig. 4a, median number of amino acids 588 vs. 432; Fig. 4b, median 5 vs. 4 domains). Notably, most IpA-generated truncated proteins still retain functional protein domains, suggesting that IpA may contribute to the diversification of the transcriptome (Fig. 4b, median 2 domains). IpA genes preferentially encode proteins with RNA-or DNA-binding or protein-protein interaction (PPI) domains, but they avoid membrane proteins. Proteins encoded by IpA genes are also enriched in repeated domains (**Supplementary Fig. 2a**, Fig. 4c), and in a majority of these we observed that IpA results in partial loss of the repeated domains. For example, the full-length protein encoded by *NFKBID* has six ankyrin domains, while the IpA-generated truncated protein retains four of them. Similarly, the full-length protein of the transcription factor PATZ1 has seven zinc finger domains, while different IpA isoforms are predicted to result in proteins with either four or five zinc fingers (Fig. 4d). The partial loss of DNA-binding domains has the potential to change DNA-binding specificity and therefore the set of target genes regulated by transcription factors. Similarly, the partial loss of PPIs can change the binding affinity to interaction partners of a protein. Taken together, these observations support the hypothesis that IpA contributes to diversification of the transcriptome and proteome.

**Figure 4:**
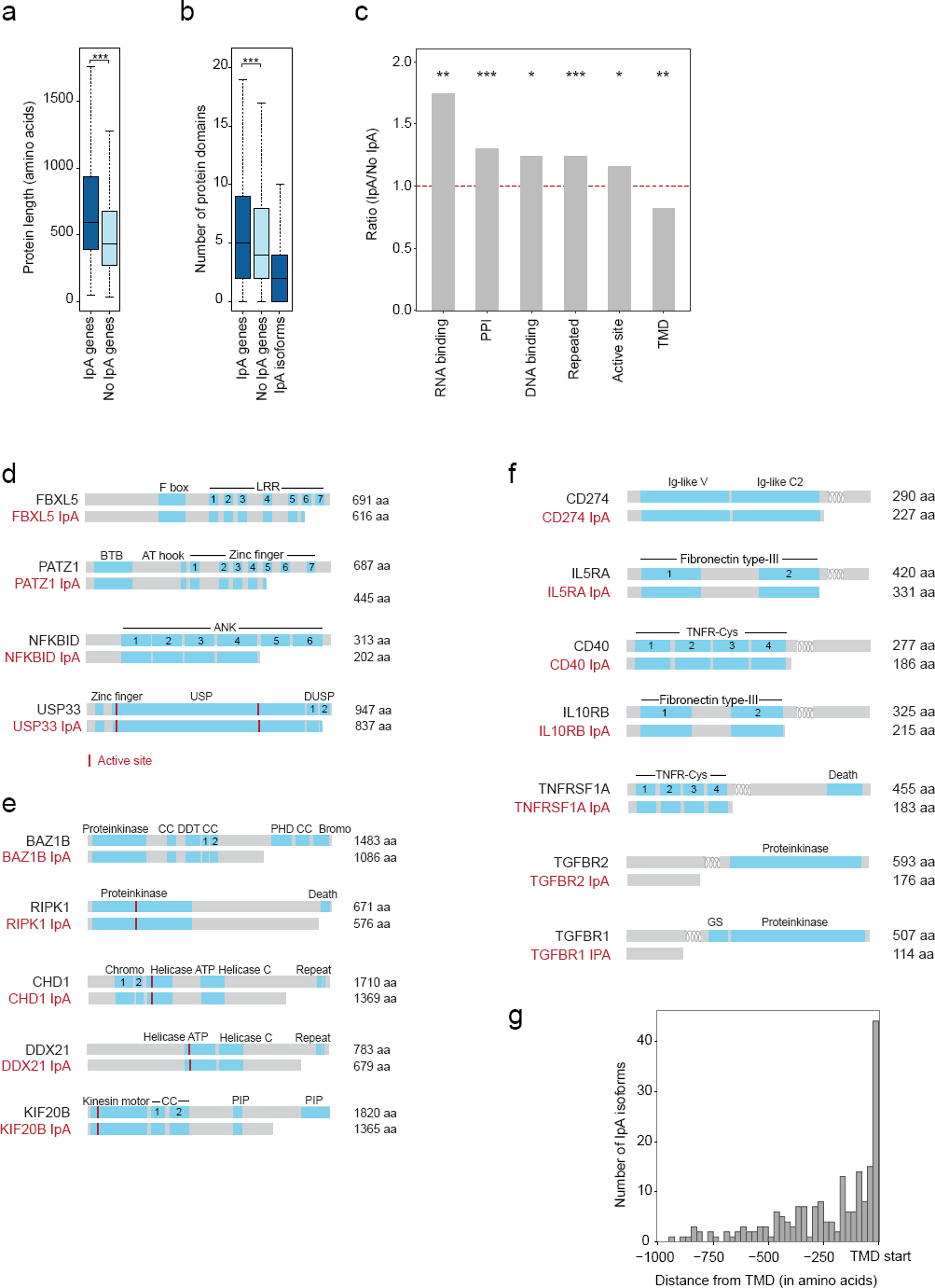
IpA isoforms diversify the proteome. a)Genes that express IpA isoforms encode significantly larger proteins compared to genes that only express full-length mRNAs (one-sided Wilcoxon rank-sum test, *p* < 10^−118^).b)Genes that express IpA isoforms encode proteins with significantly more protein domains than genes that only express full-length mRNAs (one sided Wilcoxon rank-sum test, *p* < 10^−14^). IpA isoforms retain a median of two domains.c)Genes that express IpA isoforms are enriched in proteins encoding RNA-and DNA-binding, PPI repeated domains and active sites compared to genes that only express full-length mRNAs. However, IpA genes are depleted for proteins encoding transmembrane domains (TMDs).d)Protein models of full-length and IpA-generated truncated proteins are shown in grey for examples that contain repeated domains. Known protein domains are shown as blue boxes and repeated domains are numbered.e)As in (d), but shown for enzymes that retain their active sites but lose PPI domains.f)As in (d), but shown for plasma membrane proteins. The TMD is indicated by the loops.g)Distance between the IpA event and the start of the TMD in IpA isoforms that completely lose their TMDs (n = 272).

We next investigated if there are specific protein domains that are preferentially lost or retained through IpA. Within the group of genes with a single IpA event and whose IpA isoform retains least one protein domain (n = 1,405), we found that IpA results in a preferential loss of DNA-binding or PPI domains but avoids the loss of active sites (**Supplementary Fig. 2b, c** and see **Methods**). Active sites of enzymes are the regions where substrate binding and catalysis take place. Loss of an active site would make an enzyme dysfunctional, but IpA appears to avoid this outcome. IpA genes encode diverse proteins with enzymatic functions, including protein kinases, DNA or RNA helicases or motor proteins, as shown in Fig. 4e. While the active sites are retained in the IpA-generated truncated proteins, these enzymes lose PPI domains, which may enable the enzyme to participate in different protein complexes or to change the substrate. For example, the protein kinase RIPK1 exists as full-length protein containing a C-terminal death domain, which is not included in RIPK1 IpA (Fig. 4e). BAZ1B, also called WSTF, is a multi-functional protein that contains an N-terminal protein kinase domain but has the option to also include C-terminal located coiled-coil, zinc finger and bromodomains. Also, helicases, including DDX21, DDX49 and DHX15, as well as motor proteins such as KIF20B retain their enzymatic function but generate proteins lacking interaction domains.

Membrane proteins are characterized by the presence of transmembrane domains (TMDs) and are significantly depleted among IpA genes (Fig. 4c, **Supplementary Fig. 2a**). However, we still found 673 IpA isoforms from 499 genes that encode transmembrane proteins and retain at least one protein domain. Among them, 207 IpA isoforms from 152 genes completely retained their TMDs, whereas 220 IpA isoforms from 175 genes lost their TMDs. Interestingly, IpA isoforms that retain the TMDs often encode intracellular membrane proteins that localize to mitochondria. In contrast, IpA isoforms that lose their TMDs are significantly enriched in signal peptides that are predominantly present in plasma membrane proteins (FDR, *p* < 9.1 × 10^−29^, Fig. 4f, see **Methods**). Many of them encode cytokine receptors, integrins, or growth factor receptors. Notably, regardless of the position of the TMD, the truncated protein generated by IpA usually terminates immediately before the TMD (Fig. 4g). As all of these candidates contain signal peptides at the N-terminus, the IpA isoform produces a secreted form of the cytokine or growth factor receptor.

### 5’IpA can produce robustly expressed non-coding RNAs

A large fraction of IpA isoforms that are differentially used in at least one pairwise comparison among normal immune cell types are in fact 5’IpA isoforms (487 out of 1,281). Through *de novo* RNA-seq assembly, we were able to resolve the transcript structure for 954 of the 5’IpA isoforms in our atlas (see **Methods**). Using the transcript structure we found that 469 of these have low predicted coding potential with open reading frames that are predicted to encode less than 100 amino acids^25^. Therefore, they are likely to either generate micropeptides or represent non-coding RNAs (Fig. 5, **Supplementary Fig. 3**)^26,27,28,29^. To assess potential functional consequences of expression of non-coding transcripts, we examined if RNA-binding proteins may preferentially bind to the exonized intronic sequences upstream of the IpA cleavage site. As can be seen for the examples shown in Fig. 5, the new exons are enriched for CLIP-seq peaks for RNA-binding proteins such as FUS, ELAVL1, PUM2, TAF15, and TIAL1 (binomial Z > 10, see **Methods**), which are typically enriched in the 3’UTRs of coding transcripts. As the newly exonized intronic sequences did not bind RNA-binding proteins usually bound to introns, our analysis supports the exonic nature of the predicted noncoding transcripts.

**Figure 5:**
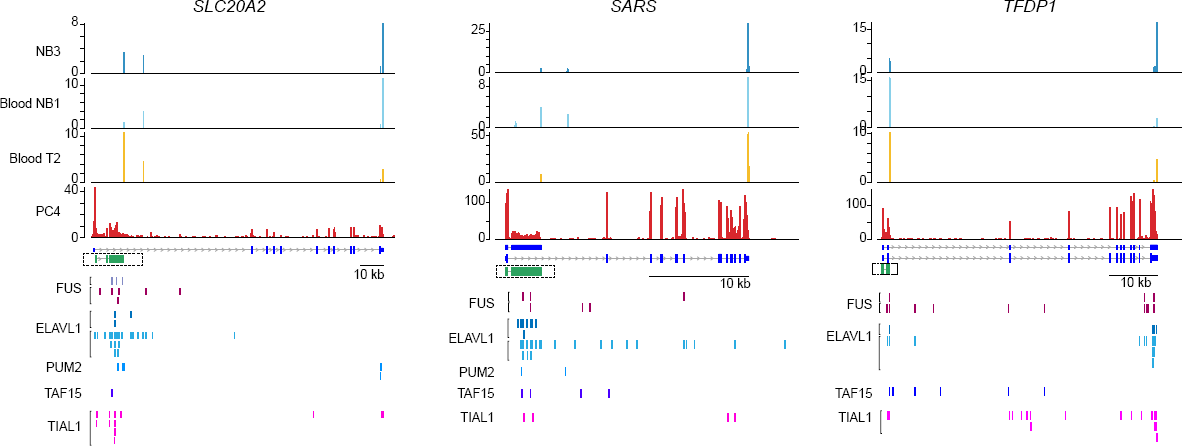
5’IpA isoforms potentially express non-coding RNAs. Examples of 5’IpA isoforms are shown as in Fig. 1b. Also shown is the structure of the assembled IpA isoform transcripts in green. Enrichment of CLIP-seq tags over exonized introns of IpA isoforms are shown for the RNA-binding proteins FUS, ELAVL1, PUM2, TAF15 and TIAL1.

### Multiple myeloma displays a widespread loss of plasma cell IpA isoforms

As alternative 3’UTR isoform expression can be altered in cancer cells ^6,30,31^, we investigated whether IpA is also dysregulated in cancer. Since PCs express the highest number of IpA isoforms among the tissue-derived B cells, we compared IpA isoform expression between normal and malignant PCs, derived from MM patients (n = 15). As MM is a heterogeneous disease, we used hierarchical clustering based on IpA isoform expression to define three patient subgroups (**Supplementary Fig. 4 and Table 2**). We then performed GLM modeling as described above to determine the differential relative expression of IpA isoforms versus full-length isoforms for each MM group compared to normal PCs. Whereas one patient group had an IpA profile comparable to normal PCs, two MM patient groups showed widespread loss of usage of PC IpA events (groups 1 and 2, Fig. 6a). We found that 44% of all PC-expressed IpA isoforms (480/1088) are lost in at least one patient group, while only 15 IpA sites show increased usage (FDR-adjusted *p* < 0.05). The significant events in patient group 1 largely represent a superset of those in group 2 (Fig. 6b).

**Figure 6:**
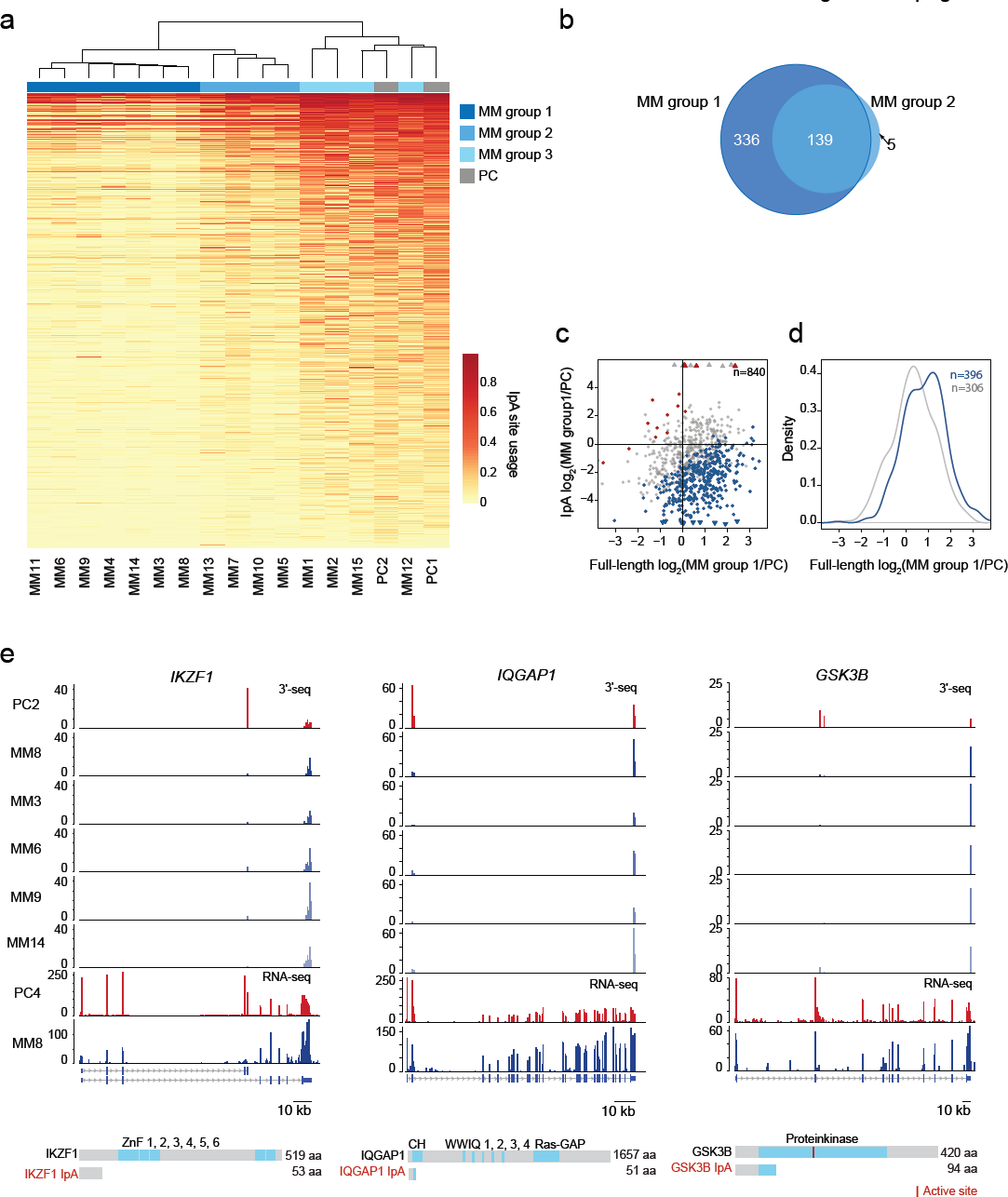
Loss of usage of IpA sites in MM. a)Heatmap shown as in Fig. 3b, but for PCs and MM patient samples. MM samples were grouped according to IpA site usage and color-coded, and the IpA isoforms with significantly different usage compared to PCs are shown (FDR-adjusted *p* < 0.05, lower usage of IpA sites in MM, n = 480; higher usage of IpA sites in MM, n = 15, not shown).b)Overlap of significantly lower used IpA sites in MM group 1 and group 2. MM group 3 is not shown as IpA site usage was very similar to PCs and only one IpA isoform was differentially used.c)As in Fig. 3d, but for MM group 1 versus PCs. Full-length and IpA isoform expression is shown and significantly different IpA isoforms are color-coded (FDR-adjusted *p* < 0.05).d)As in Fig. 3e. Shown is a significant upregulation of full-length mRNA isoform expression (one-sided KS test, *p* < 10^−8^) of genes highlighted in blue in (c).e)Examples of IpA isoforms expressed in PCs, but significantly decreased in MM samples. Shown as in Fig. 1b and Fig. 4e.

Loss of IpA isoform expression in patient group 1 resulted in a significant increase of full-length mRNA expression (Fig. 6c, d). As with differential IpA site usage between normal cell types, the genes that display differential relative IpA isoform expression in MM versus PC are enriched for annotations such as bromodomain, transcriptional regulation, and ubiquitin-like conjugation pathway (data not shown). In the majority of patient samples profiled (11 out of 15), the MM transcriptome is characterized by the loss of 480 IpA isoforms that are normally expressed in PCs. This is in contrast to 3’UTR regulation, where we found shortening of 3’UTRs in 126 and lengthening of 3’UTRs in 215 multi-UTR genes (MM group 1; data not shown).

Interestingly, one of the genes that displays loss of IpA isoform expression is the transcription factor *IKZF1,* a key gene in MM biology and also the target of lenalidomide, a recent MM therapeutic (Fig. 6e)^32^. The IpA isoform of *IKZF1* results in the loss of all zinc finger domains encoded by the full-length isoform, potentially leading to expression of a dysfunctional truncated protein isoform as it only contains 53 amino acids and no known domain. While the IpA isoform is the dominant isoform in PCs, with only minimal expression of the full-length transcript, in MM group 1 patients, expression of the IpA isoform is almost completely lost, and the full-length transcript is instead aberrantly expressed. Similarly, the gene encoding *IQGAP1,* a GTPase-activating scaffold protein that plays a role in cell proliferation in MM, largely loses expression of its IpA isoform in MM^33^. This isoform, which lacks its Ras-GTP domain as well as most functional domains, is either non-coding or at best produces a truncated protein with only a fraction of the N-terminal calponin homology domain, an actin-binding domain (Fig. 6e). Finally, in PCs, *GSK3B* predominantly expresses an IpA isoform that truncates the kinase domain and loses the catalytic site. In MM, full-length transcript expression is restored, presumably rescuing GSK3B-mediated signaling. This may contribute to MM biology as GSK3B functions as a pro-survival factor in MM^34,35,36^

## Discussion

We used 3’-seq and RNA-seq analyses to demonstrate the widespread expression of IpA isoforms across human tissues. IpA isoform expression has been observed and experimentally validated previously but has primarily been viewed as a form of alternative splicing involving alternative last exon usage^15,37^. However, as the defining event is the usage of an intronic alternative polyA signal, we instead use the term IpA. We performed comprehensive IpA analyses using 46 samples and identified 4,927 high-confidence IpA events. The majority of the IpA events described here have not been annotated thus far. Our study showed that IpA is unexpectedly widespread and especially common among normal human immune cells. As IpA isoforms are often highly expressed, they represent a normal component of the expression program in human cells and are therefore not ‘cryptic’ or ‘transcriptional noise’. Instead, their expression is regulated across normal cells and dysregulated in cancer.

The widespread nature of IpA isoform expression has escaped attention thus far, as RNA-seq analysis alone is unable to accurately identify mRNA 3’ ends. However, combining 3’-seq and RNA-seq analyses enabled us to resolve transcript structure and to identify hundreds of new non-coding RNAs as well as truncated mRNAs that are predicted to generate proteins with alternative C-termini. The IpA-generated truncated mRNAs are expressed at the same order of magnitude as full-length mRNAs and are not subject to degradation by nonsense mediated decay, since their stop codons are not premature as they are followed by conventional mRNA 3’ ends.

It is currently thought that the expression of IpA isoforms is regulated by a competition between splicing and cleavage-polyadenylation reactions^10,16^. Consistent with this model, we found that IpA genes have distinct structural and sequence properties compared to genes that generate only full-length transcripts that may predispose them toward IpA recognition. In particular, IpA genes have longer introns, longer transcription units, an enrichment of polyA signals, and a depletion of U1 snRNP signals relative to genes that only express full-length transcripts^21,22^. Nevertheless, the tissue-specific differential expression of many IpA isoforms also suggests more complex regulation of the production or stability of these transcripts. It is possible that degradation factors, including components of the RNA exosome, are downregulated in immune cells, resulting in a more frequent occurrence of IpA^38^. Alternatively, there may be an expression change of splicing factors, such as hnRNP C and U2AF65, which have been associated with the regulation of Alu exonization, one of the mechanisms known to control the expression of intronic exons^39,40^. The fact that more prevalent use of IpA signals, shorter IpA isoforms, and shorter 3’UTRs are all correlated across tissues suggests that the abundance of the same global co-transcriptional factors may be partially responsible for all three properties.

The finding that long introns and long transcription units are more susceptible to IpA suggests that the processivity of the co-transcriptional machinery may also play a role in IpA expression. Intron retention has also been observed as a prevalent feature of blood cell transcriptomes^41,42^. If we view splicing and cleavage-polyadenylation as competing processes carried out by different co-transcriptional complexes, the tendency to retain certain introns through intron retention may provide the polyadenylation machinery time to recognize IpA signals for cleavage and 3’ end processing. Interestingly, we did find a statistically significant co-occurrence of introns with IpA events and retained introns (**Supplementary Fig. 5a**). In particular, prevalence of intron retention correlates with prevalence of IpA across cell types (**Supplementary Fig. 5b**) and retained introns are enriched for IpA in each cell type examined (**Supplementary Fig. 5c**). However, IpA events in introns with no evidence of intron retention have higher IpA site usage than those in retained introns (**Supplementary Fig. 5d**). Therefore, it is unclear from our data whether intron retention is a necessary mRNA processing step prior to IpA recognition and 3’ end formation, or whether IpA recognition can occur independently of intron recognition.

One of the surprising findings of our study was an enrichment of IpA isoforms located at the 5’ end of the transcription unit that predominantly occur in immune cells. We observed 378 robustly expressed IpA isoforms that occurred in introns in 5’UTRs and thus are predicted to be non-coding transcripts. Using less stringent criteria and allowing the generation of 100 amino acids, we identified 469 of 5’IpA isoforms. These IpA isoforms are either non-coding or represent a novel source of micropeptides^26,27,28,29^. The cellular function of non-coding RNAs generated through IpA is unclear. There are reports of promoter-associated RNAs that initiate upstream of transcription start sites and that regulate transcript expression through RNA interference or interaction with epigenetic modifying enzymes^43,44,45,46^. However, in our matching RNA-seq data, we did not find read evidence upstream of transcription start sites indicating that the predicted non-coding RNAs that we observed in our study originate at the annotated transcription start sites. Additionally, we demonstrated through analysis of CLIP sequencing data that the exonized intronic sequence of the 5’IpA isoforms contains binding sites for RBPs; potentially, these non-coding RNAs serve as scaffolds for RBPs and thereby exert a regulatory role in *trans* on other RNAs.

The vast majority of IpA isoforms (n = 2,667), however, are predicted to generate truncated proteins that retain at least one domain. Notably, IpA genes encode larger proteins that contain significantly more domains than proteins generated from non-IpA genes. As the IpA-generated truncated proteins still retain a median number of two domains, the majority of proteins have the potential to be functional, thus suggesting that IpA isoforms are an important source of diversification of the transcriptome. This is supported by a significant enrichment of repeated protein domains among the proteins encoded by IpA genes. Strikingly, in the majority of cases, the repeated domains are only partially lost, thus modulating but not losing overall protein function. IpA-induced transcriptome diversification is also supported by the finding that the active sites of enzymes are retained in IpA-generated truncated proteins. Again in these cases, IpA results in proteins with similar function as the full-length proteins but different affinity or different binding partners. This suggests that the cell-type specific expression of truncated proteins generated through IpA may be a widely used mechanism to diversify the proteome, not a peculiarity of a few well-known examples like IgM.

IpA can also generate C-terminal truncations of membrane proteins. Although IpA genes avoid membrane proteins overall, IpA can mimic ectodomain shedding in transmembrane proteins. For example, metalloproteinases such as ADAM10 or ADAM17 are known to release the ectodomains of several surface receptors, including TNFα, L-selectin, TGFα or CD40^47,48^. For several of these molecules, the soluble ligands act as agonists or antagonists of the membrane-bound ligands. In particular, while membrane-bound Fas ligand kills T lymphocytes, soluble Fas ligand blocks this activity^49^. In the vast majority of the investigated cases, proteolytic cleavage occurs close to the plasma membrane, cutting at a site near the TMD, thus releasing the extracellular domain of the growth factor receptor or cytokine. Intriguingly, we found that IpA is another mechanism to produce soluble versions of membrane-bound receptors – very similar to the soluble ligands generated through proteolytic cleavage – as IpA – generated truncations also occur close to the TMD. This demonstrates that developmental regulation of membrane-bound versus secreted molecules – which was first described for IgM – is widespread and can mimic proteolytic cleavage carried out by proteases. The soluble proteins generated through IpA include CD274 IPA, encoding a truncated PD-L1. PD-L1 is an important immune checkpoint receptor ligand, and serum levels of soluble PD-L1 correlate with poor prognosis in B cell lymphomas^50^. IpA also generates a soluble tumor necrosis factor receptor 1 (TNFR1), which has been shown to block TNF activity and is associated with multiple sclerosis^51^.

As IpA isoform expression changes in different stages of B cell development as well as after environmental changes, the balance of membrane-bound versus soluble forms can change for many cytokine-or surface receptors. Importantly, IpA isoform expression is not only dynamic during development but is also dysregulated in cancer. A majority of the MM patients that we profiled showed a striking loss of IpA isoforms normally expressed in PCs. Thus, it seems that hundreds of genes avoid the generation of full-length proteins in PCs and the majority of them are re-expressed in MM samples. One gene that displays a switch-like loss of IpA isoform expression and rescue of full-length transcript expression in MM is *IKZF1,* the gene encoding IKAROS, a transcription factor and chromatin remodeler and a key therapeutic target in this malignancy^32^. Another gene with biological relevance for MM is GSK3B which has pro-survival activity in MM^34,36^. Whereas PCs express the full-length mRNA as well as the IpA isoform of GSK3B, MM samples exclusively express the full-length mRNA. As a group, genes that lose IpA expression in MM samples compared to PCs also upregulate full-length transcript expression, presumably rescuing the function of the full-length protein to varying degrees. Interestingly, not all cancer cells show depletion of IpA isoforms as we found increased rather than reduced IpA isoform expression in another B cell malignancy, chronic lymphocytic leukemia, which is presented elsewhere.

## Methods

### 3’-seq computational analyses

Preprocessing of 3’-seq libraries, read alignment (hg19) ^52^, identification and quantification of peaks were performed as described by Lianoglou et al. (2013) ^1^. The peaks were assigned to genes using RefSeq annotations. To obtain an atlas of robust cleavage events in 3’UTRs and introns, we started with all the peaks that were detected by peak calling of all the pooled samples and then followed a series of steps to filter lowly expressed peaks and the ones that potentially originate from different artifacts.

#### Removing artifacts

The peaks potentially resulting from different artifacts were identified and removed: i) peaks overlapping blacklisted regions of human genome (n = 2,841; 0.16%) (https://sites.google.com/site/anshulkundaje/projects/blacklists) ^53^; ii) internally primed peaks (Lianoglou et al. 2013) (n = 662,562; 36%); and iii) antisense peaks (n = 289,340; 16%).

#### Removing the immunoglobin peaks

Our dataset included plasma cells which are fully differentiated B cells that secrete antibodies. As plasmas cells produce massive quantities of antibodies, a large fraction of 3’-seq reads mapped to immunoglobulin loci on chromosomes 2 and 14. It was essential to account for this skewed expression of specific genomic regions in order to get a reasonable quantification for the expression of other genes. Thus, peaks (n = 11) overlapping with parts of the genome coding for immunoglobulins were removed. Even after this correction, one sample of plasma cells had a high number of intergenic reads (PC2). Thus, this sample was not used for identification of robustly expressed isoforms but only to quantify them. The library size was reduced accordingly for plasma cell samples, since the peaks described above result either from sequencing artifacts or from skewed expression of specific genomic regions.

#### Removing ambiguous peaks

Some genes in the genome overlap with each other. In such cases, it is difficult to assign 3’-seq reads to the genes accurately, and thus such genes (n = 336) were removed from further analysis. This resulted in the removal of 8,437 peaks from the atlas. Since we were interested in investigating the IpA isoforms of protein coding genes, peaks falling in introns that potentially originated from microRNAs, small nucleolar RNAs and retrotransposons were also removed (n = 4,722). Genes that were on the opposite strand but had a 3’UTR end in the intron (100 nt) of a convergent gene can create artifactual antisense peaks in the intron. Thus, peaks in introns that were close to the end of an opposite strand 3’UTR were also removed (n = 2,091). This corresponded to discarding peaks in the introns of 630 genes. There are genes where the end of the 3’UTR might fall in the intron of the downstream gene on the same strand. This would also create peaks in introns that are contributed by the preceding gene. Therefore, peaks in the intron that were within 5000 nt of the 3’ end of the 3’UTR of the previous gene were discarded (n = 2,079); the discarded peaks came from the introns of 785 genes.

#### Identification of robust isoforms

The expression levels of IpA and 3’UTR ApA isoforms were quantified by tags per million (TPM) falling in 3’-seq peaks, i.e. the read count of the peak regions was normalized by the library size of the respective sample. A gene can have many cleavage events with adequate expression levels. To examine cleavage events that represented one of the major isoforms with respect to all isoforms with a 3’ end in a given gene, these isoforms were filtered by usage. Usage is a statistic that gives an estimate of the relative expression of the isoform. As different 3’UTR ApA isoforms create the same protein irrespective of the 3’UTR length, the usage of IpA isoforms was calculated with respect to the total expression of 3’UTR ApA isoforms. IpA isoforms that end in different introns result in distinct protein isoforms (when translated), therefore, their usage was calculated relative to the total expression of both IpA isoforms and 3’UTR isoforms.

As we were interested in analyzing functionally relevant isoforms, we filtered for robustly expressed isoforms by imposing TPM and usage cutoffs. For the 3’UTR ApA isoforms, an isoform that was expressed with at least 3 TPM and with usage of 0.1 or more became part of the atlas. To focus on the most confident IpA isoforms, an IpA isoform was considered to be robustly expressed only when it was expressed with 5 TPM or more and had 0.1 usage in at least one sample. The interquartile range of the start position of the reads was also required to be 5 or more for the peaks in that particular sample to be defined as a real IpA isoform to eliminate peaks originating from PCR duplicates. These criteria helped to filter the lowly expressed isoforms as well as any possible known artifacts. Filtering for these expression criteria shrunk the atlas from 410,404 peaks to 46,923. As we were interested in IpA and 3’UTR ApA isoforms that would have different functional consequences, peaks that were within 200 nt were clustered to represent a single 3’ cleavage event. Clustering reduced the number of peaks to 40,105. After following the steps above, the atlas comprised 27,927 peaks in 15,670 genes for cleavage events of the 3’ UTRs and 3’ ends of 5,957 IpA isoforms in 3,945 genes. For downstream analysis, we only focused on the IpA isoforms (n = 5,670) of protein coding genes (n = 3,768).

### Validation and independent read evidence of IpA isoforms

We tried to corroborate the robustly expressed IpA events described thus far (n = 5,670) with external sources of evidence as described below:

#### 1. External annotation

As annotated, we consider mRNA isoforms present in RefSeq, UCSC or Ensembl. Last exons of all the existing transcripts of the hg19 annotation for Refseq, UCSC and Ensembl were obtained. These last exons were resized to include a region 100 nt downstream of the annotated end. If the 3’ end of the IpA isoform detected by our 3’-seq analysis overlapped with an expanded last exon, then it was considered to be substantiated by an external annotation. 39.52% (n = 2,241) of all the IpA events fell in the vicinity of annotated 3’ end (using the previous definition) based on an external annotation.

#### 2. RNA-seq GLM

RNA-seq read coverage is expected only over the exons and not over the introns, since the splicing machinery splices out introns during co-transcriptional processing of the pre-mRNA. However, if there is an IpA isoform that ends in an intron, then there should be RNA-seq read coverage before the 3’ end of the IpA isoform and no read coverage after the 3’ end (Fig. 1d). To test whether the upstream read coverage was significantly higher than the downstream read coverage, two windows of 100 nt separated by 51 nt upstream and downstream of the IpA 3’ end were defined. These two windows served as replicate bin counts for upstream and downstream coverage. As this was done within each single RNA-seq sample, library size normalization was not required (i.e. the size factor was set as 1 for every comparison). Significant differential expression upstream vs. downstream using DESeq ^17^ was then tested (FDR-adjusted *p* < 0.1). Not all IpA isoforms could be tested by DESeq. IpA isoforms where the defined windows overlapped with an annotated exon were excluded from further analysis. In total, 4,802 events were tested. As a control for this analysis, random introns of expressed genes that did not contain 3’ end peaks were sampled and analyzed as described above. DEseq analysis returned *p* values consistent with the null hypothesis (**Supplementary Fig. 1a**). The RNA-seq validation was applied over all the RNA-seq samples. If an IpA event was validated in any sample, then it was considered be supported by RNA-seq data. Of all the IpA isoforms, 29% (n = 1,670) could be validated by this approach.

#### 3. Other 3’-end sequencing protocols

If IpA isoforms detected by our 3’-seq protocol were also found by other 3’-end sequencing methods ^18^, we include the IpA event in our atlas of high confidence IpA events. This led to the inclusion of 1,332 IpA isoforms. The peaks reported by Gruber et al. (2016) were resized to be 75 nt width (25 nt upstream of the original start and 50 nt downstream of the original start). Overall 70% (n = 3,999) of IpA events were supported by other 3’-end sequencing protocols.

#### 4. Untemplated adenosines from RNA-seq reads (RNA-seq, polyA reads)

In RNA-seq data, some reads may overlap the 3’ end of the templated transcript and the start of the polyA tail; these reads contain untemplated adenosines and thus fail to map to the genome. Reads that did not map to the human genome were therefore used to get additional support for the IpA 3’ ends. To make sure that these reads were at the 3’ end, the reads ending with 4 or more As were trimmed. Only reads that were greater than 21 nt in length after trimming were retained. Unmapped reads from all RNA-seq samples were trimmed, and all reads with untemplated As were pooled. These reads were then aligned to the human genome. Using the aligned BAM file, all the reads that were possible PCR duplicates were further filtered out. The uniquely mappable reads that overlapped with the IpA peak (20 nt extended upstream and downstream) were counted. If an IpA isoform was supported by four or more trimmed RNA-seq polyA reads together with the presence of one of the polyA signals (AAUAAA and its variants)^54^, then the IpA isoform was considered to be corroborated by polyA RNA-seq reads.

#### 5. Highly expressed IpA isoforms

Since many of our cell types have not been previously assayed by other 3’-end sequencing methods and are also not represented well in existing RNA-seq data sets, we rescued highly expressed cell type-specific IpA isoforms by using a stringent expression cutoff (10 TPM and 0.1 usage). We also required the presence of an upstream polyA signal (AAUAAA and its variants)^54^. This step enabled us to include 323 IpA events in the atlas of highly confident IpA events.

### Expression cut-offs used for IpA and full-length mRNA expression

A gene is considered to be expressed if either the IpA isoform (≥ 5 TPM) or the full-length isoform (≥ 5.5 TPM) were expressed in 75% of the samples of the particular cell type.

### Conservation analysis

We obtained phastCons 46-way conservation scores ^20^ for 200 nts upstream and downstream of the 3’ ends of IpA isoforms to compare the mean conservation score of the 3’ ends of IpA isoforms against random introns containing polyA signals, but without IpA site usage. The random introns (n = 5,000) were chosen from IpA genes but we selected introns without IpA events, but with at least one polyA signal (AAUAAA). One of these polyA signals was randomly selected, and we obtained the phastCons 46-way conservation score for 200 nts upstream and downstream of this polyA signal.

### Identification of differentially used IpA sites

IpA site usage was calculated as the fraction of reads that map to the IpA site compared to all the reads that map to the 3’UTR of each gene. This translates into the relative expression of the truncated protein compared with the full-length protein. To identify the statistically significant changes in the usage of pA signals, we used a GLM, where we model the read counts of all isoforms across conditions by negative binomial distributions and we test for the significance of an interaction term between isoform and condition. This form of modeling approach was adapted from DEXSeq, which is formulated for testing the differential usage of exons ^23^. If a gene has multiple IpA isoforms, then the relative expression of each IpA isoform as well as the pooled full-length mRNA expression were tested independently, since different IpA isoforms are translated into different protein isoforms.

### Gene ontology enrichment analysis

Functional annotation enrichment was performed on the genes with significant differential usage of IpA sites (Fig. 3a) using DAVID with the expressed genes as the background ^24^ Functional annotation enrichment by DAVID was also performed for the genes that loose TMDs and retain TMDs with all the genes that have TMDs expressing IpA isoforms as background.

### Protein domain analysis

The information about protein domains was obtained from the UCSC UniProt annotation table (spAnnot) via the Bioconductor package-rtracklayer. Only the domains with annotation type ‘active site’, ‘domain’, ‘transmembrane region’, ‘repeat’, ‘zinc finger region’, ‘compositionally biased region’, ‘DNA-binding region’, ‘region of interest’, ‘lipid moiety-binding region’, ‘short sequence motif’, ‘calcium-binding region’, ‘nucleotide phosphate-binding region’, ‘metal ion-binding site’ and ‘topological domain’ from UniProt were used for analysis. These domains were further categorized into more broad categories: i) Active site – active site and catalytic sites, ii) DNA-binding domains – C2H2-type, PHD-type, C3H1-type, KRAB, Bromo, Chromo, DNA-binding, C4-type, CHCR, A.T hook, bZIP, bHLH, CCHC-type, CHCH, Bromodomain-like, CH1, C6-type, A.T hook-like, C4H2-type and CHHC-type, iii) Protein-protein interaction domains (PPI)-WD, ANK, TPR, LRR, HEAT, Sushi, EF-hand, ARM, PDZ, PH, SH3, RING-type, LIM zinc-binding, WW, SH2, BTB, FERM, CH, Rod, Coil 1A, MH2, WD40-like repeat, t-SNARE coiled-coil homology, Coil 1B, Cbl-PTB, Coil, CARD, SH2-like, DED, IRS-type PTB, SP-RING-type, EF-hand-like, RING-CH-type, v-SNARE coiled-coil homology, Arm domain, LIM protein-binding, GYF, PDZ domain-binding and PDZD11-binding. Also, if a region in protein was annotated with ‘Interaction with’ then that region was considered a PPI domain, iv) RNA-binding domains – RRM, SAM, KH, DRBM, RBD, Piwi, PAZ, S1 motif, Pumilio and THUMP, v) Transmembrane domains (TMDs)-transmembrane region, ABC transmembrane type-1, ABC transporter and ABC transmembrane type-2, and, vi) Repeated – Any domains that were repeated in the protein were considered repeated domains. If a gene had multiple protein isoforms, then the longest isoform was used in the analysis. The protein lengths were obtained from http://www.uniprot.org/ for *Homo sapiens.*

### Distance of IpA from TMDs

IpA isoforms for which there was positional information about the start of first TMD and those that retained at least one domain were used for this analysis. Further, we focused on IpA isoforms that completely lost all their TMDs due to the cleavage event in the intron. The distance of the retained CDR (in amino acids) by IpA from the first TMD was determined as: (Upstream CDR from IpA – Upstream CDR from the intron before the first TMD)/3.

*De novo* transcript assembly

### De novo transcript assembly

The complete transcript structure was obtained through the following steps. i) We used StringTie, an improved method for more accurate *de novo* assembly of transcripts from RNA-seq data ^55^. *De novo* assembly was performed on every RNA-seq sample with default settings using the hg19 RefSeq annotation. ii) Transcripts from multiple assemblies were subsequently unified using CuffCompare, which removes redundant transcripts and provides a set of unique transcript structures ^56^. iii) For each individual gene, we obtained the transcripts that overlapped the gene’s coordinates. We gave preference to multi-exon transcripts over single exon transcripts. For single exon transcripts, we allowed the start/end to be within 100 nt of the TSS. We gave this advantage to the single exon transcripts because the direction of transcription for these transcripts is not certain. iv) Finally, using the 3’ ends of IpA isoforms (from our 3’-seq data), we assigned transcripts with the nearest ends to these IpA isoforms.

Firstly, we identified transcripts that ended within 50 nt of 3’-seq events. If there were several assembled transcripts meeting this criterion, we chose the transcript that had the maximum number of exons. If there was a tie in the number of exons, then we chose the transcript that started closest to annotated TSS. For the remaining 3’-seq events, we assigned the nearest ending transcript. Finally, using the above defined criteria for selecting transcript structures, we determined which IpA isoforms corresponded to these assembled transcripts. If the 3’ end of the IpA isoform was within 500 nt of the defined transcript end, then we assumed that this particular transcript represented the full structure of the IpA isoform. For some IpA isoforms we observed usage of different polyA signals within the same intron. Thus, to account for such cases, for the IpA events that did not fall within 500 nt of a transcript end, we determined if it overlapped a transcript that ended within 5000 nt. If this was the case, then we assigned this transcript to that 3’ end. We were able to define the transcript architecture for n = 954 IpA isoforms (both annotated and unannotated). If the transcripts ends differed from the IpA 3’-seq events, then we defined the 3’ end determined from 3’-seq to be the real end. This was done as 3’-seq identifies 3’ ends of polyadenylated mRNAs at single nucleotide resolution and thus is more accurate than transcript ends obtained from short read assembly.

*Coding potential prediction*-To determine the probability that the 5’IpA events represented non-coding transcripts, we made use of CPAT, a tool that predicts the coding potential of the transcript based on four sequence features: open reading frame size, open reading frame coverage, Fickett TESTCODE statistic, and hexamer usage bias ^25^. For our analysis, we considered non-coding IpA isoforms to be the ones that had coding potential probability less than 0.3, had retained coding sequence less than 25%, and had ORF ≤ 300 nt (n = 469).

### Binding site enrichment of RNA-binding proteins in exonized introns

We used available CLIP (cross-linking immunoprecipitation) sequencing data of RNA-binding proteins from dorina ^57^. As the majority of CLIP studies was performed in HEK293 cells, we we focused on non-coding IpA isoforms expressed in HEK293 and only included IpA isoforms in the analysis whose exonized intron was larger than 50 nt (IpA isoforms = 62, genes = 58).

We determined if the exonized part of the intron was enriched for binding sites of RNA-binding proteins compared to other regions (introns, coding exons, 3’UTRs) of the transcription units, called ‘background’ here. We calculated the expected number of binding sites in the exonized introns using each background and compared it to the observed number of binding sites in the exonized introns. This enabled us to calculate a binomial Z-score of each CLIP experiment and each background region. We observed enrichment of binding sites of RNA-binding proteins in the exonized introns compared with introns and coding exons but no enrichment compared to 3’UTRs. The RNA-binding proteins with Z-scores ≥ 10 compared to introns or coding exons are PUM2, FUS, ELAVL1, TIAL1 and TAF15.

### Identification of retained introns

Retained introns were identified using a modified version of the IRFinder algorithm ^41^. To avoid genes with a complex genomic architecture, we removed genes that overlap with other genes in either the sense of antisense strand. An intron was categorized as retained if it satisfied the following criteria. i) There should be at least three reads spanning both the upstream and downstream exon-intron junction. ii) At least 50% of the intron length should be covered by 3 or more unique reads. Mappability of introns could be a limitation in this case, and thus we focused only on introns that had at least 50% uniquely mappable sequence relative to its complete length. iii) To ensure adequate expression of the flanking exons, the median coverage over the flanking exons was required to be 10 reads or more. iv) Since the introns should have more coverage than background noise, we considered introns to be retained if the ratio of median coverage over the intron to median coverage of the upstream and downstream exons was at least 10%.

An intron was annotated as retained if it fulfilled all criteria in at least 66% of the RNA-seq samples of a given cell type. Introns retained in 33% or fewer samples were flagged as not retained while the introns that were retained in more than 33% samples but less than 66% of RNA-seq samples were removed from the analysis. For a 3’ end of an IpA isoform to occur in a particular intron, the intron must contain a polyA signal or one of its variants ^54^.

Our data showed that some genes had very high coverage over almost all the introns of the gene, presumably due to sequencing artifacts. We determined the (median coverage over all the introns)/(median coverage over all the exons), and if this ratio was ≥ 0.2 then these genes were flagged for removal.

The number of introns that would have IpA and IR simultaneously by chance were calculated as – Probabilty of IpA ×Probabilty of IR × Number of expressed introns with polyA signal

